# RSV Downregulates IL-21/IL-21R on TFH cells via PD-L1 induction in APCS impairing protective humoral responses

**DOI:** 10.1101/203133

**Authors:** Rodrigo Benedetti Gassen, Tiago Fazolo, Deise Nascimento de Freitas, Thiago Borges, Fábio Maito, Daniel A. G. Bueno Mendes, André Báfica, Luiz Carlos Rodrigues, Ana Paula Duarte de Souza, Cristina Bonorino

## Abstract

Respiratory syncytial virus (RSV) is the major cause of hospitalization for children under two years of age. RSV vaccines are currently unavailable, and children suffering from multiple reinfections by the same viral strain, fail to develop protective memory responses. Follicular helper T (TFH) cells specialize in providing B cell help to antibody production and affinity maturation, mainly via IL-21 secretion. Although RSV-specific antibodies can be detected upon infection, how they are generated and their relevance against disease protection has not been fully examined. Here, we observed that RSV expands a functionally impaired murine TFH cell population *in vitro* and *vivo*, with downregulated IL-21R expression and IL-21 production. IL-21 treatment of RSV-infected mice, however, increased TFH cells frequency, enhanced the germinal center reaction and improved protective humoral immune responses by increasing viral protein F specific antibody avidity and neutralization capacity. *In vivo*, it protected from RSV infection, decreasing lung inflammation. Passive immunization with purified IgG from IL-21 treated RSV-infected mice protected against RSV infection. Both viable and UV-inactivated RSV induced PD-L1 expression on B cells and DCs, however, only in DCs a direct effect of RSV was detected. Blocking PD-L1 during infection recovered IL-21R expression in TFH and B cells and increased secretion of IL-21 by TFH cells in a DC-dependent manner. Our results unveil a novel pathway by which RSV affects TFH cells activity, reducing levels of IL-21 and its receptor, by increasing PD-L1 expression on APCs. These results highlight the PD-L1/IL-21 axis importance for the generation of protective responses to RSV infection.

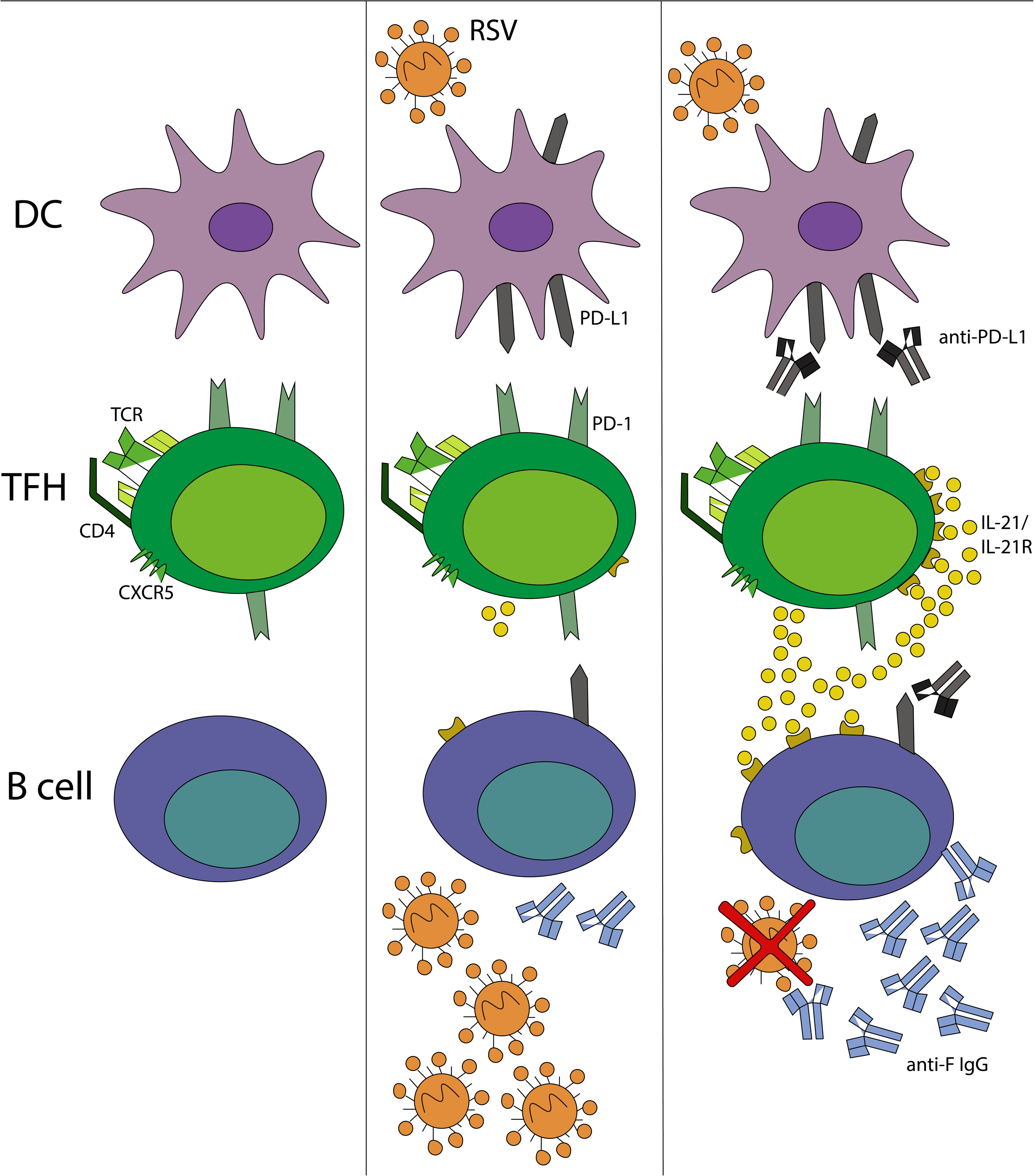
GRAPHICAL ABSTRACT. RSV infection impairing IL-21 secretion by TFH cells via PD-L1induction in dendritic cells and B cells. Low levels of IL-21 lead to poor RSV-specific humoral immune responses, low antibody titer, avidity and neutralization capacity. PD-L1 blockade can upregulate IL-21 secretion, and IL-21 treatment restores the entire immune humoral responses, resulting in protection against RSV infection.

## INTRODUCTION

Respiratory syncytial virus (RSV) is the leading cause of lower respiratory tract infection in infants, responsible for about three million hospitalizations and 66,000 deaths every year in children under two years of age (1). RSV-infected infants develop upper respiratory tract symptoms, often progressing to bronchiolitis and/or pneumonia, and even respiratory failure, which frequently needs mechanical ventilation and intensive care unit therapy (2). There is no effective vaccine available against RSV, and passive immunization with monoclonal antibodies is used only in high risk infants, in which RSV infection can be deadly. However, it is not only in premature children that RSV infection may be problematic or deadly; infants, elderly and pregnant women are also target populations (3), reinforcing the need for development of an effective vaccine.

In humans, neutralizing RSV-specifics antibodies are formed in upper respiratory tract, however, re-infection with the same RSV strain is frequent in healthy, immune competent individuals (4,5). The protective relevance of serum antibody levels remains unclear (6,7). Serum antibody response was loosely correlated with protection; however, there was a correlation of nasal preexisting RSV-specific humoral response with resistance to re-infection (8). Other studies have correlated the presence of serum high avidity RSV-specific IgG with protection (9); although these antibodies have a short life in children (10) and adults (11). Nevertheless, RSV morbidity and mortality are mainly associated with 2-4 months of age infants when titers of maternal antibodies are decreasing and have not yet been replaced by an endogenous antibody responses (12). In mice, the RSV antibody response induced by formalin-inactivated RSV (FI-RSV) is non-protective (13). Formalin inactivation results in conformational changes of the RSV fusion protein (14), which is required for the generation of effective neutralizing antibody responses (15). Moreover, antibodies induced by FI-RSV are low affinity due to poor toll-like receptor (TLR) stimulation (16).

Effective B cells responses require help from follicular helper T (TFH) cells. TFH cells are predominantly found in germinal centers (GC) of secondary lymphoid organs (17–19) and produce high levels of IL-21. This cytokine is known to increase the affinity of antibodies and induce immunoglobulin class switching (20,21). IL-21 acts on naive B cells in conjunction with co-stimulatory signals which drive differentiation of either GC B cells or plasma cells (22). The specific provision of TFH cell-derived helper signals to GC B cells is proposed to be the major driver of antibody affinity maturation (23,24). Therefore, we hypothesized that RSV modulates TFH cells, preventing B cell help, targeting humoral immune response efficiency.

By employing a murine model of RSV infection, our results unravel an immunological mechanism by which RSV modulates TFH cells and GC reactions. We found that RSV can induce PD-L1 (Programed Death-ligand 1) in dendritic cells (DCs) and B cells, decreasing B and TFH cells functions. This correlated with a decreased ability of TFH cells to produce IL-21 and downregulation of IL-21R, leading to low avidity RSV-specific humoral immune response. Treatment with recombinant IL-21 reduced RSV-mediated impairment of GC and TFH cell functions as well as improved animal survival. Our results underline the importance of the PD-1/PD-L1 pathway and IL-21 adjuvant activity in the generation of effective anti-RSV protective antibody responses.

## METHODS

### Viruses and Cells

Vero cells (ATCC CCL81) were cultured in Dulbeccol’s modified Eagle’s medium (DMEM, Gibco-BRL) containing 10% fetal calf serum (FCS, Gibco-BRL) and gentamicin (0.08 mg/ml, NOVAFARMA), maintained at 37°C with 5% of CO_2_ and used to propagate RSV A2 strain and HSV-1 KOS strain. Viral plaque-forming units (PFU) were identified using an RSV F protein-specific antibody (Millipore, Billerica, MA, USA). Lytic plaques assay was used to calculate PFU of HSV-1. Inactivation of the viruses was performed at UV light for 30 min, at room temperature.

### Animals

Female BALB/c mice ranging from 6 to 8 weeks old were purchased from the Biological Center of Experimental Models (CEMBE) of PUCRS. Mice were housed in CEMBE facility with water and food *ad libitum*. All animal procedures were performed in accordance with the guidelines of the Federation of Brazilian Societies for Experimental Biology and approved by the Ethics Committee for the Use of Animals of PUCRS (CEUA-PUCRS; protocol number #13/00341).

### Infection and treatment

For *in vitro* infection, splenocytes were isolated after lysis of red blood cells. Cells were seeded at 5×10^5^ cells per well in a 96 well-plate and were stimulated either with 10^2^ PFU/ml of HSV-1 or RSV or the same virus inactivated by UV for 30 min. As negative controls, uninfected Vero cells were used and processed similarly to RSV infected cells. Alternatively, cells were treated with 0.5 µg of anti-PD-L1 antibody (clone MHI5, eBioscience) or control IgG (BioXcell) 1 hour before infection. The supernatant was collected at 12, 24, 48, 72 and 96 hours post-infection for analysis. After four days, cells were labeled with specific antibodies and analyzed by flow cytometry.

For sorting, splenocytes were isolated after lysis of red blood cells and stained with specific antibodies for DCs (CD11c^+^ CD19^−^), B cells (CD19^+^CD11c) and TFH cells (CD4^+^CXCR5^+^). Cells were co-cultured (5×10^3^ DCs, 3×10^4^ TFH cells or 5×10^4^ B cell per well) in different combinations, and later infected with 10^2^ PFU/ml of RSV for 4 days. Alternatively, cells were treated with 0.5 µg of anti-PD-L1 antibody (clone MHI5, eBioscience) or IgG (BioXcell), 1 hour before infection, and analyzed for IL-21R and PD-L1 expression.

For *in vivo* infection, mice were divided into 5 groups: two groups were infected intranasally with 10^7^ PFU of RSV, and one of this group received subcutaneous treatment with 0.5 µg of recombinant IL-21 (eBioscience) diluted in PBS. Two groups received PBS intranasally, and one of this group received treatment with recombinant IL-21 subcutaneously. And the last group was infected with 10^7^ PFU of HSV-1 via intraperitoneal. IL-21 was administered on days: 4, 8, 14 and 18 post-infection. The blood collection occurred on days 0, 4, 8, 14, 18 and 21 post-infection. Mice were euthanized at day 21 post-infection and spleens and lungs were harvested for further analysis.

### ELISA

IL-21 concentration in supernatant and serum was determined by capture ELISA (R&D Systems), according to manufacturer’s instructions.

For the quantification of IgG RSV specific antibodies on mice serum, 96-well plates were sensitized overnight with RSV F protein (Sino Biological Inc.), blocked for 1 hour with blotto (5% milk, 0.05% tween in PBS 1X buffer) and serum was added in dilutions of 1/10, 1/100, 1/500, 1/1000, 1/5000 and incubated for 2 hours at room temperature. Rabbit anti-mouse antibody HRP-conjugated (Invitrogen) and TMB (Life Technologies) was used to development. Plates were read at 450nm in EZ Read 400 Microplate reader (Biochrom).

ELISA to measure the avidity of anti-RSV antibody was conducted in the same manner, however the plate was washed with a 6M Urea solution. The results were expressed as an Avidity Index (AI), which was calculated as previously described by Freitas et al (9). The avidity of RSV-specific total IgG was classified according to values that had been predetermined arbitrarily defined as low (<50%), intermediate (50-70%), or high (>70%).

### Neutralization assay

Antibody neutralization capacity assay was performed as previously described by Zielinska et al (26), with modifications. Four-fold serial dilutions from 1:10 to 1:10240 were prepared in virus diluent (DMEM 0% FCS and 1% gentamicin (0.08 mg/ml, NOVAFARMA). Serially diluted serum was challenged with an equal volume of the RSV-A2 strain, previously tittered to give ~150 PFU per 50 µl of inoculum. The serum/virus mixtures were incubated at 37°C, 5% CO_2_ for 1 h.

96-well plates plated with Vero cell monolayers were infected with 50 µl/well (in triplicates) of the serum/virus mixture. Plates were blotted with 0.5% methyl cellulose, prepared in DMEM with 2% FBS and incubated at 37°C, 5% CO_2_ for 3 days to allow plaque formation. To detect the syncytium formation, wells were incubated with primary anti-RSV antibodies (Millipore) and secondary antibody HRP-Rabbit anti-mouse IgG (Millipore) for 1 hour at 37^º^C. Blocking occurred with blotto (5% milk, 0.05% tween in PBS 1X buffer). For revelation, we used 4-chloro-1-napthol with 0.01% H_2_O_2_ for 20 min in dark. Syncytium was counted in optical microscope and PFUs calculated by ratio of the number of syncytium by multiplying the dilution with virus medium volume in the plate.

### Flow Cytometry

Splenocytes were labeled with: anti-IL-21R (PE, 4A9 clone) and anti-CD19 (APC-Cy7, ID3 clone) from BD Biosciences, anti-CD4 (eFluor710-PerCP, clone RM4-5), anti-PD-1 (PE-Cyanine7, clone J43), anti-Bcl-6 (PE mgl191E clone), anti-CXCR5 (APC, clone SPRCL5), anti-ICOS (FITC, clone 7E17G9), anti-CD45R (FITC, clone RA3-6B2) and anti-CD274 (PE, clone MIH5) from eBioscience, and anti-Ki67 (PE, clone 16A8) from BioLegend. For intracellular staining, we used Foxp3 staining buffer Set (eBioscience) and cells were analyzed using a FACSCanto II (BD Biosciences) with the FACSDiva software (BD Biosciences). Alternatively, for sorter cells were used FASCAria (BD Biosciences).

### Histology and Immunohistochemistry

Lungs and spleens were fixed with 10% formalin, embedded in paraffin and cut into 5 μm-thick sections. Hematoxylin and eosin staining was performed in slide sections to evaluated inflammation scores. For immunohistochemistry, slides sections were deparaffinized with xylol and endogenous peroxidase activity was blocked by incubation with 3% H_2_O_2_ in methanol. Antigen unmask was performed by incubating the slides in 0.1 mol/L citrate buffer, pH 6, for 30 min at 95°C, followed by cooling at room temperature for 1h. Sections were blocked in PBS with 4% bovine serum albumin (BSA), 5% mouse serum, and incubated with primary antibody anti-PD-L1 (clone MIH5, eBioscience) at 1:1000 dilution in PBS, 1% BSA, and 1.25% mouse serum. Biotinylated goat-derived secondary antibodies were detected by the avidin-biotin-horseradish peroxidase complex method (Dako Systems) using 3,3-diaminobenzidine-tetrahydrochloride (Sigma-Aldrich) as a substrate.

### Real Time-PCR

Total RNA was extracted from the lungs of infected animals using Viral RNA/DNA Mini Kit (PureLink® - Invitrogen) following manufacturer’s instructions. cDNA was synthesized with random primers using Sensiscript® Reverse Transcription kit (QIAGEN®). The quality of cDNA for each sample was tested by amplification of the endogenous β-actin gene using specific primers from TaqMan Assay (Applied Biosystems). Samples that did not amplify for β-actin were excluded. Real-time PCR was performed for the amplification of the RSV F protein gene using the primers and specific probes (forward-5’-AACAGATGTAAGCAGCTCCGTTATC-3 ’, reverse-5’-GATTTTTATTGGATGCTGTACATTT-3 ’and probe 5’-FAM / TGCCATAGCATGACACAATGGCTCCT-TAMRA / −3’). For standard curve a ten-fold serial dilutions of 6×10^7^ copies of a plasmid with RSV F protein sequence were added to the same plate of qPCR in duplicate. The results were measured by StepOne™ Real-Time PCR System (Applied Biosystems) and used for further quantification of the samples viral load.

### Passive immunization

Mice were separated into 3 groups, the first group received naive IgG serum, the second group received IgG serum from RSV-infected mouse and the latter group received IgG serum from RSV-infected and IL-21 treated mouse. For IgG purification was used Protein A-Sepharose column (Sigma) following manufacturer’s instructions. Each mouse received intraperitoneal 300µg of purified IgG. After two days, animals were infected with 10^7^PFU of RSV. Mice were euthanized at day 5 post-infection and lungs were harvesedt for further analysis.

### Statistical analysis

The significance for differences between the groups was analyzed with one-way ANOVA test followed by a Bonferroni post-test or t-test were applied to parametric data using GraphPad Prism software (San Diego, CA, USA). Values demonstrated in graphs are mean and standard deviation (S.D.) and a level of significance of p < 0.05 was established for the analyses.

## RESULTS

### RSV induces TFH proliferation *in vitro*

Given the importance of TFH cells in helping B cells to produce antibodies, we analyzed whether RSV could modulate TFH cells proliferation *in vitro*. Mice splenocytes were infected with 10^2^ PFU/ml of HSV-1 or RSV for 4 days. HSV-1 was used as a positive control since it induces high levels of high affinity HSV-specific antibodies (27–29). We found that both RSV and HSV-1 increase the frequency of TFH (CD3^+^CD4^+^CXCR5^+^PD-1^+^) cells compared to uninfected controls (Fig. 1, A and B). TFH cells induced by RSV also expressed ICOS and Bcl6 (Fig. 1 C, supplementary figure 1). RSV-induced TFH cells were proliferating (as seen by Ki67 expression) compared to non-TFH (CD3^+^CD4^+^CXCR5^−^PD-1^−^) cells (Fig. 1, D and E).

**Figure 1.**
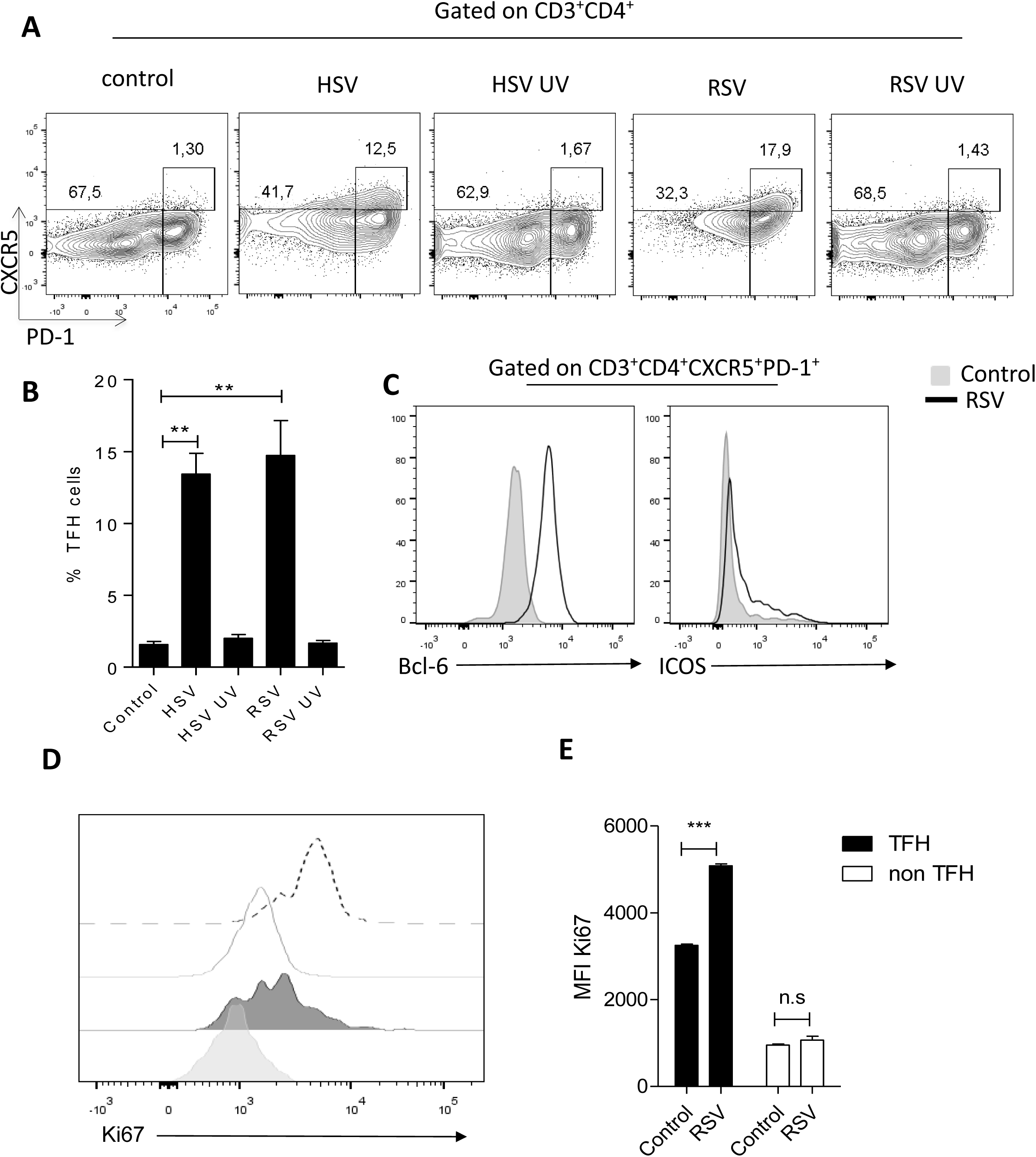
RSV induces TFH cell proliferation *in vitro*. Splenocytes were incubated with ether live RSV or HSV-1, UV inactivated RSV or HSV-1, or Vero cells supernatant for 4 days. **(A)** Representative and **(B)** Mean TFH (CD3^+^CD4^+^CXCR5^+^PD-1^+^) cells percentages. **(C)** Bcl-6 and ICOS expression by TFH cells (RSV, black line; HSV-1, black dotted line; negative control, hatched gray histogram). **(D)** Representative histograms and **(E)** MFI of Ki67 expression on TFH (Control, gray line; RSV, black dotted line) and non-TFH CD4^+^ T cells (CD3^+^CD4^+^CXCR5^−^PD-1^−^) (Control, hatched gray histogram; RSV, hatched dark gray histogram). Results are the mean of one representative experiment of three performed. *P <0.05; **P<0.01; ***P<0,001.

### RSV decreases IL-21 secretion and IL-21R expression by TFH and B cells

We next investigated whether RSV could affect the function of TFH cells. Production of IL-21 is an important indicator of TFH activity (30). IL-21 is necessary for antibody affinity maturation and B cell differentiation, and is known to upregulate the expression of its receptor on CD4^+^ T cells. No IL-21 production was detected when RSV was added to splenocyte cultures, in contrast to what was observed for HSV-1 (Fig. 2 A). *In vivo*, IL-21 serum levels were undetectable even 4 days post-infection (Fig. 2 B), again contrasting to the abundant IL-21 induction observed during HSV-1 infection. *In vitro, RSV* also reduced IL-21R expression on TFH cells (Fig. 2, C and D) and B cells (CD3^−^CD19^+^) (Fig. 2, E and F), while HSV-1 increased IL-21R expression on these cell populations (Fig. 2 C-F), suggesting the regulation of the IL-21/IL-21R axis was specific to RSV. These data indicate that RSV could negatively modulate the function of TFH cells by downregulation of IL-21R expression, as well as of IL-21 production.

**Figure 2.**
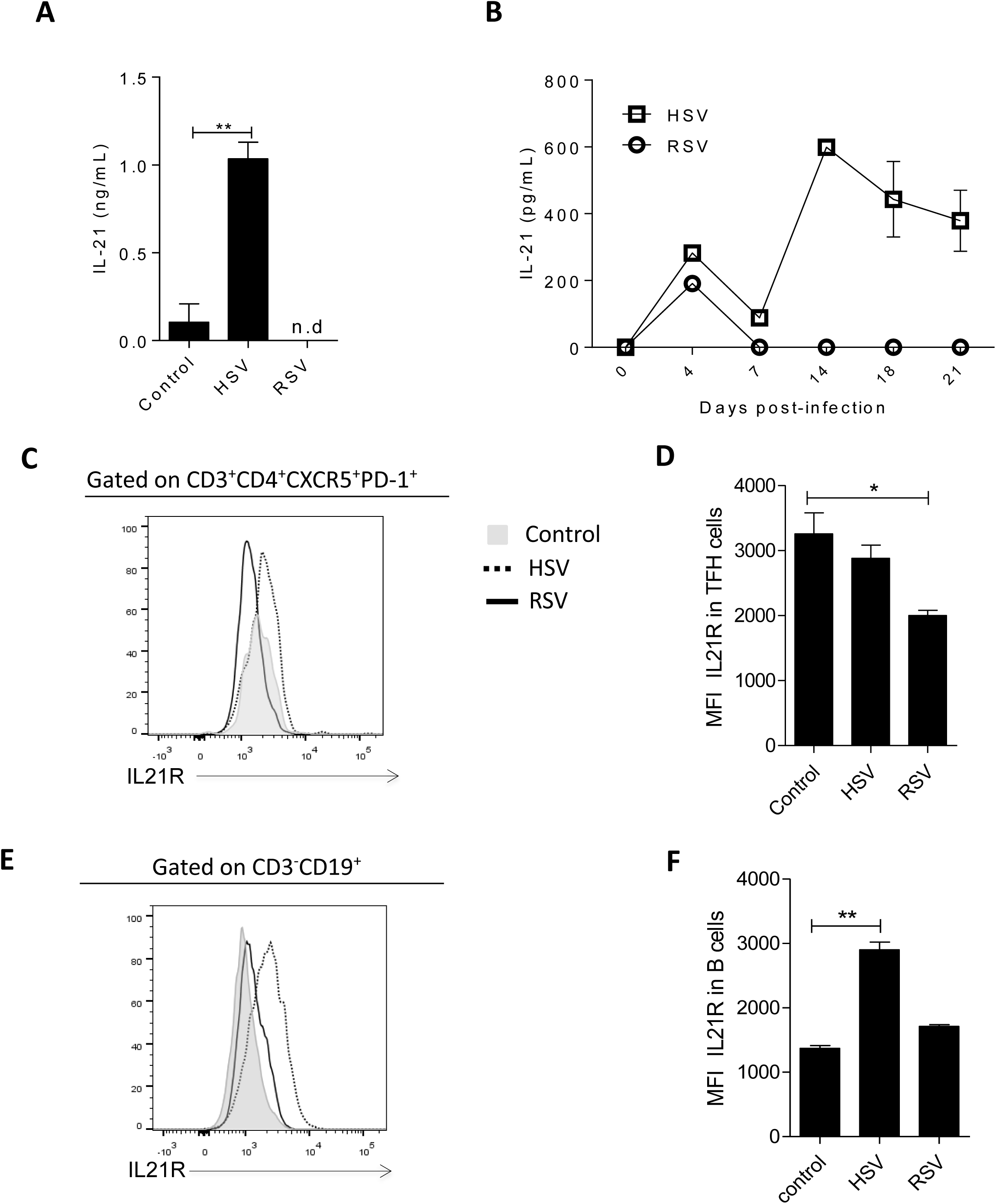
RSV infection decreases splenocyte IL-21 secretion as well as IL-21R expression in TFH and B cells. Splenocytes were incubated with ether live RSV or HSV-1, UV inactivated RSV or HSV-1, or Vero cells supernatant) for 4 days. (**A)** IL-21 in supernatant of cultured infected splenocytes. **(B)** Serum IL-21 levels in mice (three per group) infected with RSV or HSV-1, measured by ELISA. **(C)** Representative histogram and **(D)** MFI of IL-21R expression in TFH cells. **(E)** Representative histogram and **(F)** MFI of IL-21R expression in B (CD3^−^CD19^+^) cells. (RSV, black line; HSV-1, black dotted line; negative control, hatched gray histogram) Results are the mean of one representative experiment of three performed. *P <0.05; **P<0.01; ***P<0,001.

### IL-21 treatment increases B cell follicle size, IgG production, antibody avidity and neutralization capacity

We hypothesized that the low levels of IL-21 observed upon RSV infection (Fig. 2 B) would lead to decreased humoral responses against the virus. IL-21 is a key cytokine for humoral immune responses, however the influence of this cytokine in GC reaction and antibody avidity during RSV infection was so far unexplored. To evaluate the IL-21 action in RSV infection *in vivo*, we infected mice with RSV and treated them with recombinant, endotoxin-free IL-21 (Supplementary figure 2). Accordingly, we observed that animals infected with RSV showed undeveloped B cell follicles in the spleen (Fig. 3 A), differently from HSV-1 infected mice, which presented greatly augmented follicles compared to uninfected controls. This was reversed by IL-21 treatment. Exogenous IL-21 led to increase in sizes of splenic B cell follicles during RSV infection (Fig. 3 A) as well as increased frequencies of B and TFH cells (Fig 3B and C). Finally, treatment with IL-21 also improved titers (Fig. 3 D), avidity (Fig. 3 E), and neutralization capacity (Fig. 3 F) of anti-F protein specific antibodies.

**Figure 3.**
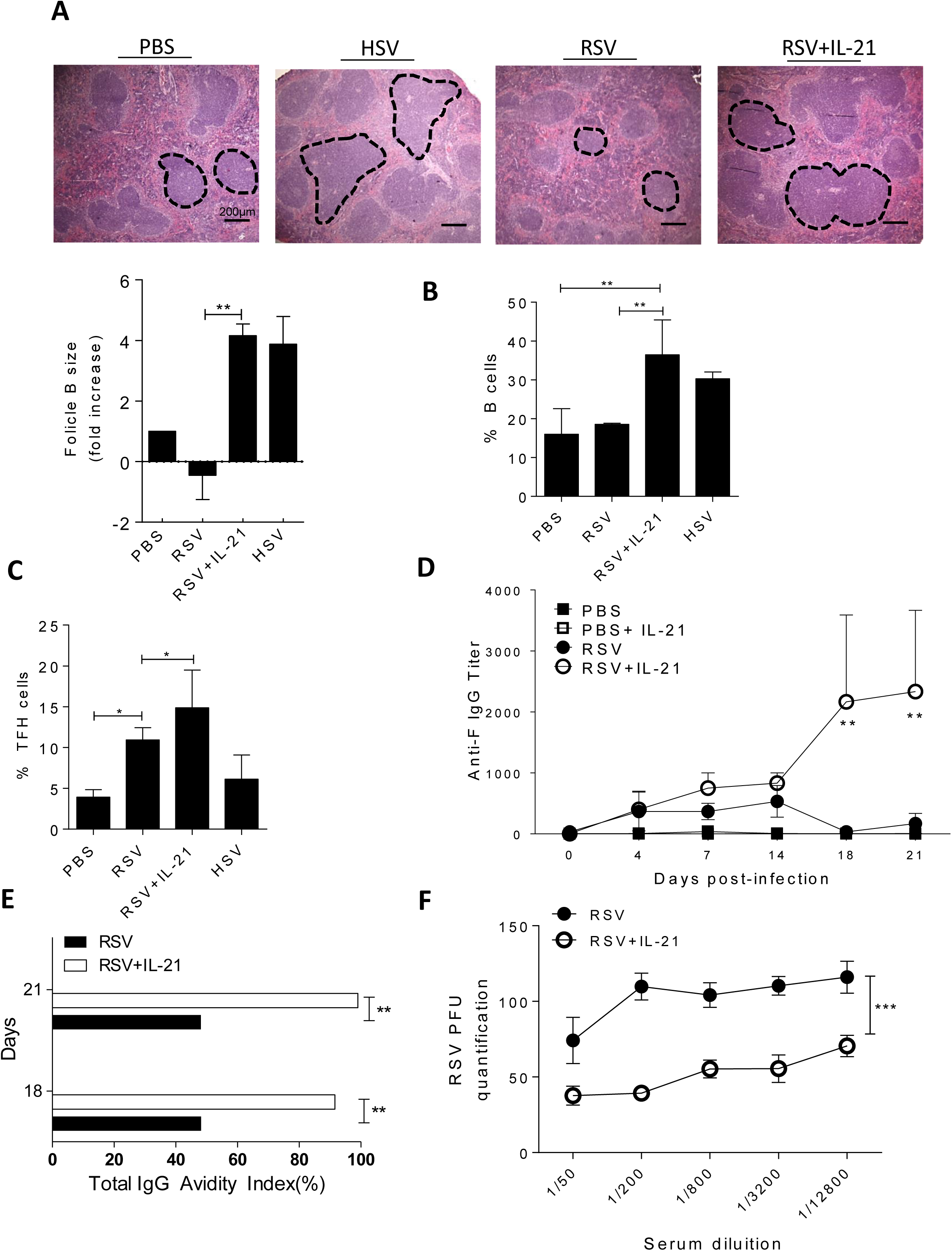
*In vivo* IL-21 treatment increases B cell follicles size, IgG production, avidity and neutralization capacity. BALB/c mice were infected with 10^7^ PFU of RSV or HSV-1 (intranasally or intraperitoneally, respectively) and treated subcutaneously with 4 doses of 0.5 µg of IL-21. **(A)** Spleen HE histology, 40x magnification. Bar scale: 200µm. Quantification of B follicle size, by fold increase. **(B)** Percentages of B cells. **(C)** Percentages of TFH cells. **(D)** Anti-F IgG titers measured by ELISA. **(E)** Total IgG avidity index on days 18 and 21 after infection. **(F)** Neutralization capacity assay. Results are the mean of one representative experiment of three performed using three mice per experiment. *P <0.05; **P<0.01; ***P<0,001.

### IL-21 treatment protects RSV infected animals and decreases lung inflammation by specific humoral immune response

To evaluate the protective roles of IL-21 *in vivo*, similar experimental design was performed (Supplementary figure 2). While pulmonary injury was observed in lungs from untreated RSV-infected mice, (Fig. 4 A, arrows show cell infiltration and reduction of bronchi lumen space), treatment with IL-21 reduced inflammation as well as improved survival compared to untreated infected mice, reducing weight loss in infected animals (Fig. 4 B and C). IL-21 treatment also decreases numbers of RSV copies in lungs in RSV-infected mice (Fig. 4 D). These results suggest that IL-21 is essential for protection from RSV infection by activating TFH and B cells function. To further investigate the immunologic mechanisms of protection, we pretreated (two days before RSV infection) mice i.p. with 300µg of purified IgG from naïve serum and RSV-infected mice serum, either treated or not with IL-21. The results, depicted in Fig. 4 E, indicate that protection can be mediated by IgG alone elicited by IL-21 treatment in infected mice. Thus, low production of IL-21 during RSV infection affect GC reactions, dramatically impairing the generation of a protective, anti-RSV humoral response.

**Figure 4.**
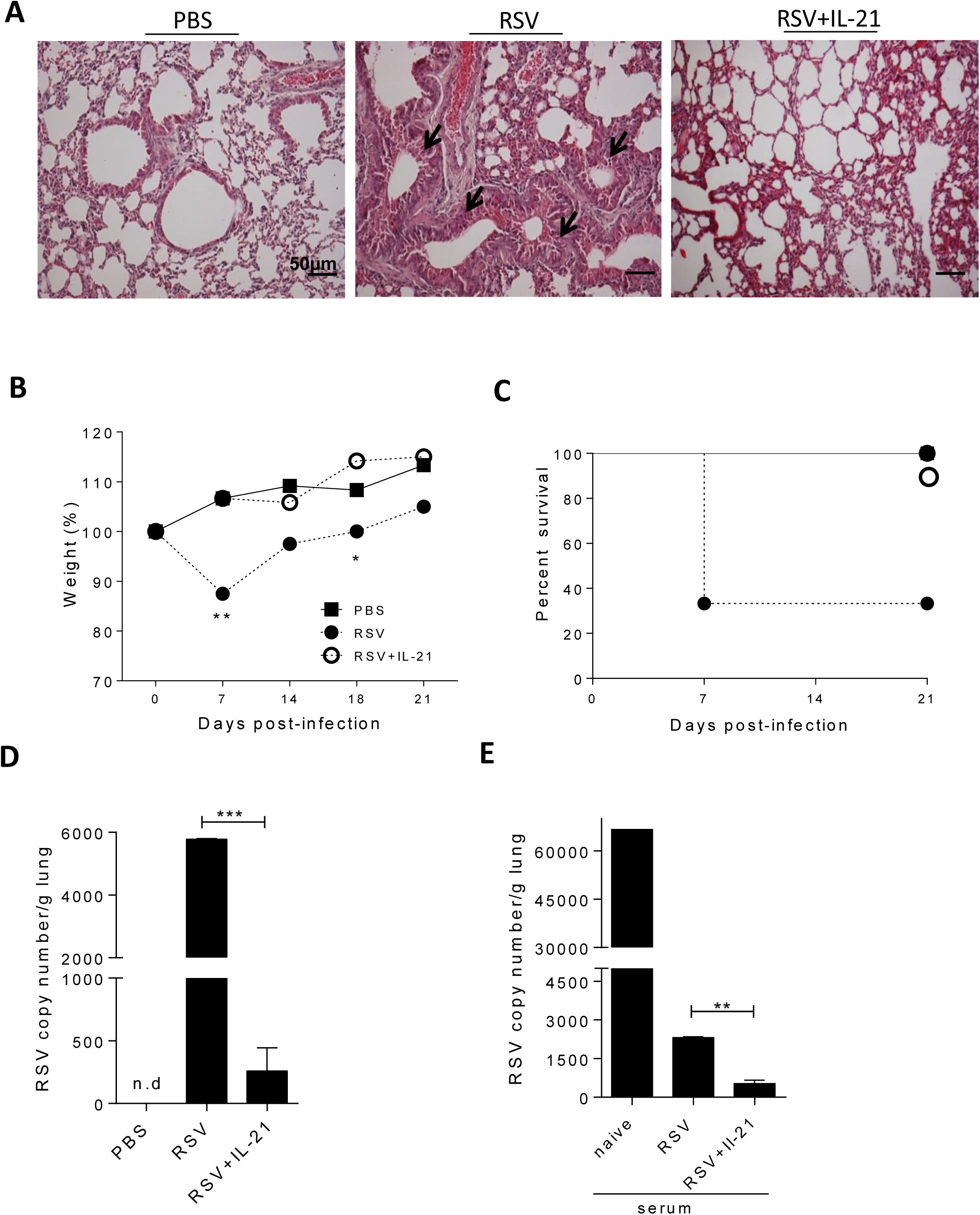
*In vivo* IL-21 treatment protects RSV infected animals and decreases lung inflammation. BALB/c mice were infected with 10^7^ PFU of RSV or HSV-1 (intranasally or intraperitoneally, respectively) and treated subcutaneously with 4 doses of 0.5 µg of IL-21. **(A)** Lung HE histology (100x magnification), 21 days post-infection. Bar scale: 50µm. Arrows indicate bronchial constriction and inflammation. **(B)** Weight loss, plotted over time. **(C)** Kaplan-Meier survival curves. **(D)** RT-PCR quantification of RSV copies in lungs, 21 days after infection. **(E)** Passive immunization with purified IgG from naïve, RSV-infected mice or IL-21 treated RSV-infected mice measured of RSV copies in lungs by RT-PCR. Results are the mean of one representative experiment of three performed using three mice per experiment. *P <0.05; **P<0.01; ***P<0,001.

### PD-L1 blockade recovers IL-21R expression on TFH and B cells and increases IL-21 secretion by TFH cells

We next investigated possible mechanisms involved in the RSV-induced downregulation of IL-21 levels. Previous studies demonstrated that RSV can upregulate PD-L1 expression on DCs (31). This leads to a decrease in humoral responses, affecting TFH viability (32). Both DCs (CD3^−^CD11c^+^) and B cells from RSV-exposed splenocytes were found to display upregulation of PD-L1 (Supplementary Fig. 3). To determine if RSV could directly induce PD-L1 upregulation in either of these cell populations, we sorted DCs and B cells, and infect them *in vitro* with RSV. Only DCs upregulated PD-L1 expression as a direct effect of RSV, and DCs also presented higher PD-L1 expression compared to B cells (Fig. 5 A and B). In vivo, RSV infection also led to upregulation of PD-L1 expression, confirmed by immunohistochemistry of infected mice spleens on day 21 post-infection (Fig. 5 C). These results led us to hypothesize that the reduction of IL-21 levels could be linked to the induction of PD-L1 in DCs following RSV infection

**Figure 5.**
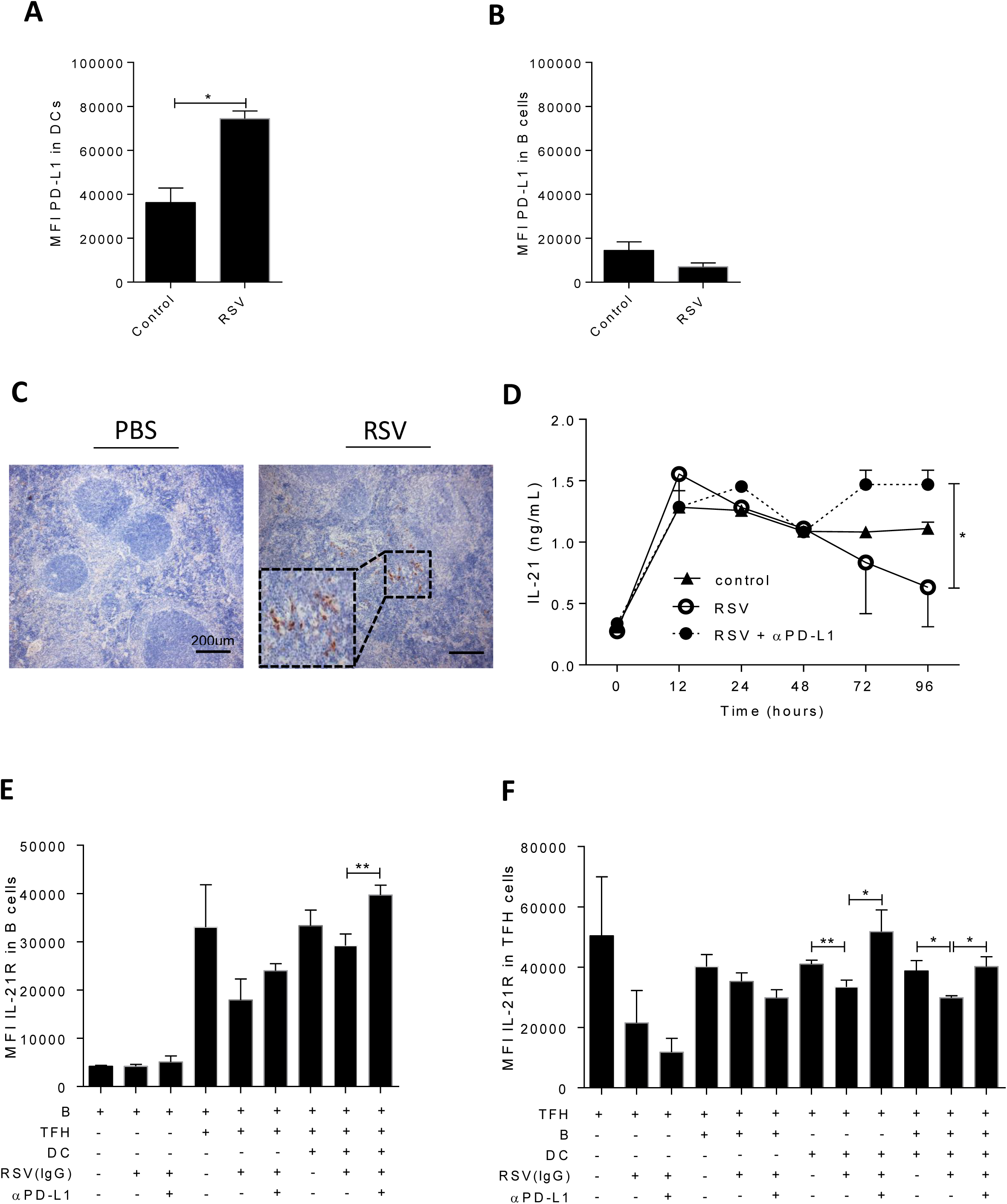
PD-L1 blockade recovers IL-21R expression in TFH and B cells and increases IL-21 secretion by TFH cells. DCs (CD11c^+^ CD19^−^) and B cells (CD11c^−^ CD19^+^) from BALB/c mice splenocytes were sorted and incubated with RSV for 4 days**. (A)** MFI of PD-L1 in sorter DCs. **(B)** MFI of PD-L1 in sorter B cells. **(C)** Spleen PD-L1 immunohistochemistry 21 days after infection (three mice per group) with 10^7^ PFUs of RSV, 40x magnification. Bar scale: 200µm. Zoom black dotted square: 100x magnification. **(D)** IL-21 levels in supernatant of infected splenocytes, with or without αPD-L1 treatment. **(E)** MFI from IL-21R in coculture sorted B cells. **(F)** MFI from IL-21R in coculture sorted TFH cells. Results are the mean of one representative experiment of three performed. *P <0.05; **P<0.01; ***P<0,001.

The prediction from our hypothesis was that PD-L1 blockade could recover the ability of TFH cells to produce IL-21 in the presence of RSV. To test that, we blocked PD-L1 *in vitro,* and infected mouse splenocytes with 10^2^ PFU/ml of RSV for 4 days. Blocking PD-L1 increased IL-21 production by RSV-infected cells, as measured by ELISA (Fig. 5 D). Two days post-infection, IL-21 levels began to decrease in supernatants. At 96h post-infection, PD-L1 blockade significantly restored IL-21 secretion (Fig. 5 D). In sorted cells, PD-L1 blockade also recovered IL-21R expression on B cells (Fig. 5 E), however only when B cells were co-cultured with TFH cells and DCs. TFH cells also recovered IL-21R expression upon PD-L1 blockade (Fig. 5 F), demonstrating that reduction of IL-21R expression is linked to PD-L1 function in a DC-dependent manner. These data suggest that engagement of the PD-L1 pathway by RSV impairs TFH secretion of IL-21. This in turn leads to downmodulation of IL-21R expression, both in TFH cells and B cells, contributing to impaired RSV-specific humoral responses (Graphical abstract).

## DISCUSSION

In this study, we reveal a mechanism by which RSV can evade effector immune responses, interfering with TFH cells function, and consequently impairing the generation of protective antibodies.

Recent studies report that impairing TFH differentiation and/or function is an important virulence mechanism for different viruses. Lymphocytic choriomeningitis virus (LCMV) negatively modulates antibody responses by killing TFH cells via NK cells cytotoxicity (34). HIV infection induces TFH cells proliferation, however recruits these cells to work as a subclinical virus reservoir during antiretroviral therapy (ART) (35). In addition, HIV infection is associated with decrease of T follicular regulatory cells (TFR) function, leading to TFH cells proliferation and increase of viral replication (36). This might be one of the reasons why HIV humoral immune responses are generally non-protective. Progressive CD4^+^ T cell loss, non-TFH, is a hallmark of chronic HIV infection, with a subsequent B cell dysfunction and poor antibody responses to vaccines (37). In RSV infections, neutralizing antibodies are formed in upper respiratory tract, however the relative contribution of anti-RSV humoral responses is still controversial. Employing several immunological in vivo and in vitro experimental models, we provide evidence that RSV inhibits TFH cells function, resulting in impaired protection.

RSV induced TFH cells proliferation, and that observation alone could indicate that RSV infection would not affect these cells. However, proliferation by itself cannot be considered as a synonym for function. IL-21 is an important indicator of TFH activity, and while IL-21 production was clearly affected by RSV infection, this was not true for HSV-1, indicating these two viruses employ different strategies regarding TFH cells.

Previous studies reported modulation of IL-21 expression as a strategy employed by pathogens to evade effector immune responses. For example, serum IL-21 levels and the frequency of IL-21-producing TFH cells in blood was lower in HCV patients compared to healthy individuals (38). Lower frequencies of IL-21–producing CD4^+^ T cells also is associated with reduced proliferation and increased expression of the inhibitory receptors like CTLA-4, Tim-3 and PD-1 on HCV-specific CD8^+^ T cells in a viral persistence state (39). In fact, IL-21 performs potent and specific effects on mucosal antiviral responses, assisting viral clearance and regulating pulmonary T and B cell responses (40). IL-21 treatment has been tested against several types of viral pathogens (HBV (41), HCV (39), HIV (42,43)). Our results demonstrated that treatment of RSV-infected mice with recombinant IL-21 had a protective effect in lung inflammation, reducing weight loss and mortality. They also corroborate results from Dodd et al, in which they demonstrated that IL-21 depletion during priming exacerbates immunopathology after RSV challenge and reduces antibody production, showing the importance of IL-21 in RSV infection (44). Studies in humans suggest that RSV can modulate IL-21 secretion. In a cohort study that characterized the primary and secondary cytokine response to RSV infection, IL-21 was not detected in swab nasal secretion samples from infants recruited during two consecutive winters (45). Our data indicated that treatment with IL-21 successfully enhanced humoral responses to RSV, leading to increases in B cell follicles, anti-F (fusion RSV protein) IgG titers, antibody avidity and neutralization capacity. High-avidity anti-RSV IgG antibodies have been shown to confer protection in infants under 3 months old (9). We observed that the antibodies generated upon IL-21 treatment in infected mice were highly protective upon transfer to naïve animals that were challenged *in vivo* by RSV. Our data agrees with reports that IL-21 is most important at the beginning of a humoral response, activating AID (Activation-induced cytidine deaminase), promoting immunoglobulin switch class and somatic hypermutation.

In addition, IL-21 increases germinal center reactions and induces IL-21R expression on activated B cells and TFH cells (46–48). IL-21R is required to generate TFH cells, GC reaction, B cells, plasma cells and plasmablasts in mice (49). We observed that IL-21R was downregulated on TFH and B cells in RSV-infected splenocytes *in vitro*, consistently with the impairment of IL-21 production. In humans, IL-21 or IL-21R defects cause severe primary immunodeficiency, reduced NK, T and B cell activity and leading to multiple infections (50). Our results indicate that downregulation of IL-21R is another important mechanism associated with the impairment of B cell responses against RSV in mice, evidencing the relevance of the IL-21/IL-21R axis in the generation of protection to this virus.

Finally, we found that decreased IL-21 production during RSV infection is dependent on PD-L1 induction. TFH cells naturally express PD-1 (17), and PD-L1, one of the classic PD-1 ligands, is responsible for immune response homeostasis, negatively modulating PD-1-expressing T cells (51). PD-1-PD-L1 interactions reduce Akt phosphorylation in PD-1 expressing cells, leading to decreases in function and cell survival (52). PD-L1 has been previously reported to be increased in DCs (31,53) and B cells (54) of mice infected with RSV. HSV-1 infection is known to induce protective specific antibodies (55), differently from RSV, suggesting that the induction of PD-L1 by RSV is one of the mechanisms associated with deficient antibody production in RSV infection. In our hands, RSV, but not HSV could directly upregulate PD-L1 expression in DCs; B cells upregulation of PD-L1 needed DCs and T cells. It is possible that the mechanism by which PD-1/PD-L1 interactions inhibit TFH cells function is by downregulation of c-Maf expression, a transcriptional factor involved in transactivation of both the promoter and enhancer of the IL-21 gene (56). Transcriptional levels of c-Maf are increased upon tolerizing immunotherapy (57) correlating with increased PD-L1 levels. Accordingly, Cubas et al demonstrated that HIV infection increased PD-L1 frequency in GC B cells leading to reduction of TFH cells proliferation and IL-21 production (58). Accordingly, several studies have suggested that blockade of PD-1 or PD-L1 may lead to a reversion of T cell dysfunction in the context of chronic infection (59). PD-L1 blockade in regulatory B cells can recover TH1 cell activity in visceral leishmaniasis in a canine model (60). When we blocked PD-L1 in vitro during RSV infection, IL-21 secretion was increased. Our results agree with Zhou et al who observed that IL-21 serum levels were inversely correlated with the high intensity of tumor PD-L1 expression in patients with hepatocarcinoma (61). A previous study demonstrated that PD-L1 expression on lung DCs controls inflammation, and anti-PD-L1 treatment of RSV-infected animals induced pro-inflammatory cytokines, exacerbating pulmonary inflammation and host disease (54). In that study, they found only a moderate enhancement of TFH cells numbers, and no improvement of antibody responses, at day 6 after infection (54). However, we found a significant increase in serum RSV-specific antibodies at day 14 post-infection in mice treated with IL-21, but not at day 6 as described by Yao et al. Thus, we believe that the benefit of anti-PD-L1 treatment on RSV antibody response could be overridden by the lung inflammation it would induce – and an early course of Il-21 therapy might constitute an interesting approach to circumvent the TFH cell function impairment that ensues upon RSV infection.

## ACKNOWLEDGEMENTS

This study was supported by grants from Conselho Nacional de Desenvolvimento Científico e Tecnológico (CNPq), Fundação de Pesquisa do Estado do Rio Grande do Sul (FAPERGS), Coordenação de Aperfeiçoamento de Pessoal de Ensino Superior (CAPES) and Pontifícia Universidade Católica do Rio Grande do Sul.

